# Rich chromatin structure prediction from Hi-C data

**DOI:** 10.1101/032953

**Authors:** Laraib Iqbal Malik, Rob Patro

## Abstract

Recent studies involving the 3-dimensional conformation of chromatin have revealed the important role it has to play in different processes within the cell. These studies have also led to the discovery of densely interacting segments of the chromosome, called topologically associating domains. The accurate identification of these domains from Hi-C interaction data is an interesting and important computational problem for which numerous methods have been proposed. Unfortunately, most existing algorithms designed to identify these domains assume that they are non-overlapping whereas there is substantial evidence to believe a nested structure exists. We present an efficient methodology to predict hierarchical chromatin domains using chromatin conformation capture data. Our method predicts domains at different resolutions and uses these to construct a hierarchy that is based on intrinsic properties of the chromatin data. The hierarchy consists of a set of non-overlapping domains, that maximize intra-domain interaction frequencies, at each level. We show that our predicted structure is highly enriched for CTCF and various other chromatin markers. We also show that large-scale domains, at multiple resolutions within our hierarchy, are conserved across cell types and species. Our software, Matryoshka, is written in C++11 and licensed under GPL v3; it is available at https://github.com/COMBINE-lab/matryoshka.

## INTRODUCTION

The 3D structure and folding of chromatin has been shown to influence many biological processes within the cell. These include cell replication, differentiation and gene expression (1, 2), as well as alterations leading to disease (3). Recent advances in chromosome conformation capture (3C) technologies (4), that combine chemical cross-linking and high-throughput sequencing, have led to the discovery of densely packed regions of chromatin referred to as topologically associating domains. These domains are found to be conserved across cell types and species, reflecting their biological importance, and their boundaries are known to be enriched for several epigenetic marks, suggesting that these domains play a role in epigenomic regulation of expression (1, 5, 6, 7).

Several methods have been developed to identify these domains using data from Hi-C — a high-throughput experimental assay that allows genome-wide conformation capture (8, 9). These methods are based on a variety of different approaches, but most focus on exploiting particular statistics and properties of contact frequencies in the resulting data. Dixon et al. (1) introduced the concept of the directionality index, which measures the difference in contact frequency upstream and downstream of a particular chromosomal locus. Treating the directionality index as a spatially-varying statistic over the chromosome, they use a Hidden Markov Model to determine a set of domain boundaries. This statistic was then employed in several other studies (10, 11). Similarly, the arrowhead algorithm, introduced by Rao et al. (12), performs a transformation on the contact matrix designed to enhance domain boundary signals. The algorithm then determines the positions of high-scoring “corners” to determine domains (thus the algorithm’s name). Instead of predicting domains directly, some methods provide change points along the diagonal of the contact frequency matrix (13, 14), but this leaves the question of which chromatin regions are *not* in domains unresolved. Filippova et al. (15) introduce a dynamic programming approach that predicts domains by maximizing a score based on normalized, intra-domain interaction frequencies. The algorithm is run at multiple resolutions and a consensus domain set is returned with the goal of predicting domains that persist across multiple scales. However, all these algorithms have an underlying assumption that chromatin domains are non-overlapping or are not nested.

There is significant evidence to believe that chromatin folding is hierarchical, wherein sub-domains combine to form larger super-domains, instead of a sequence of non-overlapping or non-nested domains. This was initially predicted in the *Drosophila* genome by Sexton et al. (6). Further studies across different cell types and species have supported this claim. Gibcus et al. (16) went on to explain the possibility of inter-domain interactions, along with the intra-domain interactions, in mammalian genomes, including mouse and human. It was shown by Filippova et al. (15) that domains predicted by their method at different size scales tend to be more nested (i.e. hierarchical) than what would be expected in a collection of appropriately randomizeddomains with the same size distribution. There is also theoretical evidence to believe that the chromatin structure is hierarchical as shown by replicating its statistical properties on a heteropolymer chain and observing the structure of the resulting folding pattern (17).

Here, we introduce a new, efficient algorithm to derive a nested hierarchy of domains from chromatin conformation capture data. Initially, our method optimizes an objective function to obtain an optimal set of non-overlapping domains at a collection of different resolutions (i.e. size scales) (15). The resolution values are then clustered based on the variation of information distance (18) between the corresponding domain sets. This clustering is used to determine discrete levels of the hierarchy. In order to obtain consensus domains at each level, we use a scoring function that is proportional to the frequency of interactions within the domain but is normalized for variation across domain sizes. We analyze the biological significance of the hierarchical domains predicted by our method, Matryoshka, in a number of ways and also compare our results against the only other publicly available tool, TADtree, for identifying hierarchical chromatin domains (19). We show that, across multiple levels of our predicted hierarchy, the boundaries of domains are statistically significantly enriched for chromatin binding factors and modifications known to be associated with domain boundaries. We also test the conservation of multiple levels of the hierarchy across cell types and species, and find that significant conservation occurs at multiple levels. Across a variety of datasets, we demonstrate that our method can efficiently determine a domain hierarchy and can automatically account for variations in nesting and domain sizes in a data-dependent manner.

## MATERIALS AND METHODS

### Algorithm Overview

In order to find the set of nested domains in Hi-C data, we designed a multi-step algorithm that aims to predict a collection of domains such that inter-domain interaction frequencies are maximized. The algorithm first predicts an optimal set of domains across a wide range of resolutions, and then clusters and nests these domains in a data-driven manner to produce a coherent hierarchy representative of the input contact matrix. The algorithm is illustrated in Figure 1, and the phases of the algorithm are explained in detail below:

**Figure 1.**
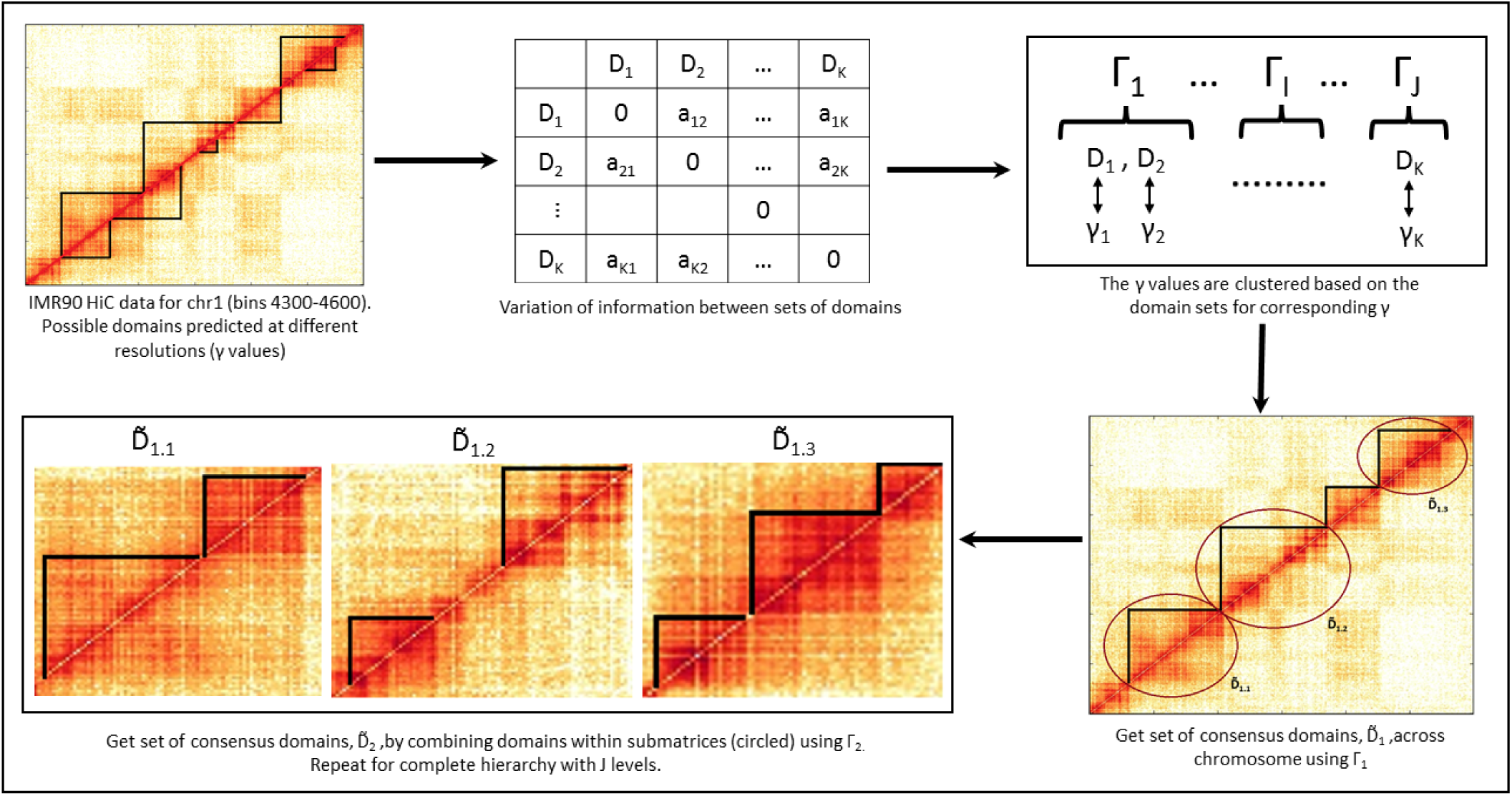
Overview of main steps of the hierarchical domain finding algorithm of Matryoshka

1. A set of non-overlapping domains is predicted at each resolution, where domain sizes tend to vary across resolutions. The heatmap in Figure 1, made using HiCPlotter (20), gives an idea of the hierarchical structure observed in HiC data.
2. The variation of information between domain sets is calculated. This is used as a distance metric for clustering the sets.
3. The domain sets are clustered and corresponding *γ* (resolution) value clusters are used for building the hierarchy of chromatin domains.
4. A set of consensus domains is obtained based on a quality score using the relevant *γ* values at each level of the hierarchy. For the first level, the set of consensus domains is across the whole matrix and for subsequent levels, submatrices predicted as domains at the higher level are used.

### Identifying putative domains across resolutions

Matryoshka takes as input an *n* × *n* interaction frequency matrix **A**, where each entry **A_ij_** represents the interaction frequency between chromosome locations (bins) *i* and *j*, and a set of resolution parameters, Γ, where each γ ∈ Γ ≥ 0. Using the method of Filippova et al. (15), a set of non-overlapping domains *D_γ_* is identified for each *γ*. Each *D_γ_* maximizes the following objective:

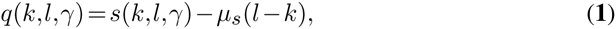
where *k* and *l* are respective genomic positions along the chromosome, and

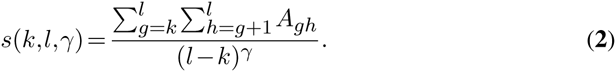

Here, *μ_s_*(*l – k*) is the mean value of *s*(*k,l,γ*) over all sub-matrices of **A** with length *l – k*. As Filippova et al. note, *γ* is inversely proportional to domain size.

In order to obtain a set of domains in the matrix, the following dynamic program is run over the length of the chromosome. This program enumerates the optimal set of domains in the sub-matrix defined by the first *l* positions on the chromosome, such that the objective function is maximized.

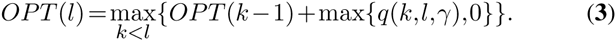

Additionally, we further filter the domains based on their boundary indices (19). This filtering, not originally applied by Filippova et al., is a reflection of the amount of shift in interaction frequencies around the boundary, and we find that it substantially reduces the number of “spurious” domains called by the algorithm. Valid boundaries should have a larger shift and, therefore, we consider domains where at least one of the boundaries has an index value greater than the mean boundary index for the whole matrix **A**. The boundary index for any position *i* is calculated as follows

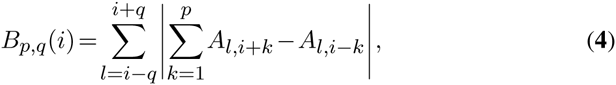
where *p* is the interval containing *i* and *q* is the length (i.e. window size) we wish to use for calculating the difference in interaction frequency upstream and downstream of *i*. Values for *p* and *q* are set to 3 and 12, respectively, as used by Weinreb et al. (19).

Given a collection Γ of resolution parameters with |Γ| = *K*, we apply this dynamic program over all *γ ∈* Γ, and obtain *K* sets of domains. The set of domains returned at each resolution are non-overlapping, but domains across resolutions may overlap. Smaller γ values result in solutions with larger domains and vice versa. These domain sets are then used to cluster similar solutions across resolutions, and the consensus domains of each cluster are used to construct the different levels of the hierarchy.

### Clustering domains to generate hierarchy

Domains obtained across resolutions from the first step are clustered based on the variation of information distance between them (18). The method for calculating variation of information between two sets of domains is described by Fillipova et al. (15). For any two domain sets, *D_i_* and *D_j_*, new derivative sets, *C_i_* and *C_j_*, are constructed such that *C_i_* contains all the domains and the inter-domain regions from *D_i_* and similarly *C_j_* is constructed from *D_j_*. The probability of seeing an interval *x_i_* = [*a_i_,b_i_*] in a derivative set for chromosome of length *L* is defined as *p_i_ =* (*b_i_ – a_i_*) /*L*. In the same way, the joint probability is defined as 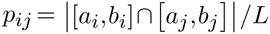. Using these probabilities, the entropy of a derivative set *C_i_* is computed as

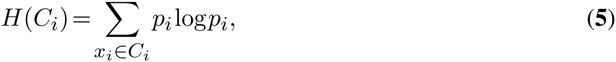
and the mutual information is computed as

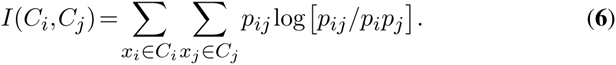

Finally, the variation of information between two sets is defined as *VI* (*C_i_, C_j_*) = *H*(*C_i_*) *+ H* (*C_j_*) *–*2*I* (*C_i_, C_j_*).

From these distances, the *K × K* variation of information matrix **V** is constructed. Each entry **V***_ij_* of this matrix provides the VI distance between the set of domains at resolutions *i* and *j*. Next, we use a clustering procedure to obtain a grouping of the *K* domain sets into a collection of *J* ≤ *K* clusters. Rather than allow clusters to consist of groups of domain sets at arbitrary resolutions, we restrict clusters to consist of collections of domain sets at contiguous values of the resolution parameter — this also allows us to employ a simple dynamic program to obtain an optimal set of clusters, by turning the clustering problem into a problem of finding an optimal partitioning of the domain sets across values of *γ*. Consider a particular partitioning of *K* domain sets into *t* disjoint intervals, given as 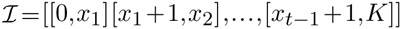s. We define a cost for this partition as

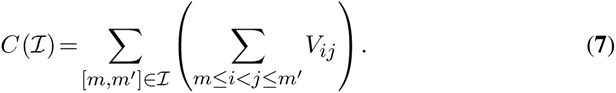

We seek a partitioning of our *K* domain sets into a collection of intervals that minimizes this cost. Given the desired number of intervals, *ℓ*, we can determine the optimal intervals by finding those that minimize the following objective:

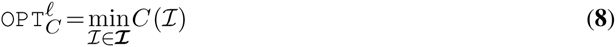

Further, this objective can be minimized efficiently via dynamic programming. Consider the objective 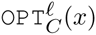, which defines the cost of an optimal set of *ℓ* intervals that cover domains at resolutions 0 through 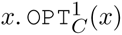 is trivial (simply the interval [0,*x*]), and

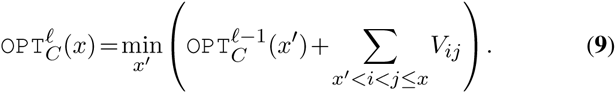

If we consider computing 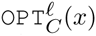 for increasing values of *x* and increasing values of *ℓ*, the optimal solution for the overall partitioning problem 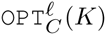 can be computed in *O* (*Kℓ^2^*) time; the actual set of intervals obtaining the optimal score can be recovered via backtracking.

The optimal number of clusters is decided based on the maximum silhouette value (21), averaged over all the points in the dataset, of the returned clustering which is defined as

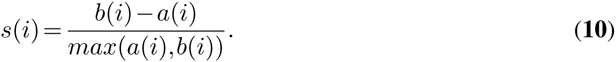

Here, *i* is a particular domain set, *a*(*i*) is the average distance of *i* from all points within the cluster and value *b*(*i*) is the lowest average distance of *i* from points in a cluster other than its own, called the neighboring cluster of *i*. The number of clusters for which the average of this value over all domain sets is maximum is chosen as the optimal cluster number for the domain sets across all resolutions. Based on this clustering, the domain sets at the corresponding *γ* values are clustered and then used to identify consensus domains at each level of the hierarchy.

### Building hierarchy using consensus domains

Given the sized *K* set of *γ* values that are split into *J* clusters, {Γ_1_, Γ_2_,…,Γ*_J_*} where 1 < J < *K* and *γ* values are sorted in ascending order, we construct a hierarchy of domains with *J* levels. Domains at any level *i* are constructed using *γ* values from within a cluster Γ*_i_*. A non-overlapping set of domains is derived at each level using a quality score independent of *γ* and domains at any level *i* completely cover the domains at any level *j > i*. Domains at coarser levels (for example level 1) are identified using larger *γ* values than those at finer levels (for example level *J*).

At level 1 of the hierarchy, a multiset of domains *D*_1_ is obtained using the interaction matrix **A** as described above. Instead of using the complete *γ* set, only the first cluster, Γ_1_, which has the largest *γ* values, is used. In order to obtain a set of non-overlapping consensus domains for level 1 of the hierarchy, the problem is reduced to the weighted interval scheduling problem (15, 22), where each domain in *D*_1_ is assigned a quality score that corresponds to its priority as follows:

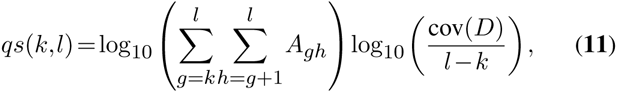
where cov(*D*) is simply the length of the chromosome covered by the complete set of domains *D* obtained in the first step.

This quality score normalizes the sum of the interaction frequency *A_gh_* between genomic loci *k* and *l*, which increases logarithmically with the domain size, against the ratio of length covered by the domain. This ratio is calculated over the complete length covered by domains in *D* instead of the chromosome length in order to disregard non-domain regions, which may cover a large portion of the chromosome. This quality score gives us the ability to compare domains of vastly different sizes across resolutions so that we can extract a set of non-overlapping consensus domains while reducing bias due to domain sizes. The result of solving the weighted interval scheduling problem with the quality scores defined above is a set of consensus domains, 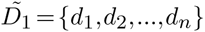, for the first level of the hierarchy.

Now, for each domain 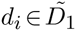, we get a submatrix of the interaction matrix **A**[*a_i_*,*b_i_*] such that the size of this submatrix is defined by the boundaries of the domain where *d_i_* = [*a_i_,b_i_*]. On this submatrix, we repeat the steps explained above to get a set of domains at different resolutions defined by the values from Γ_2_. Then we get a set of non-overlapping consensus domains using weighted interval scheduling that is placed at the second level of the hierarchy. This procedure is repeated for the all the domains within *D*_1_. In a similar way, we use {Γ_3_,Γ_4_,…,Γ*_J_*} in order to get domains at lower levels of the hierarchy, *{D_3_,D_4_,…,D_J_}* and then extract consensus domains from it for each level, 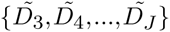, to eventually construct the complete hierarchy of chromatin domains.

### Data and testing

We have tested Matryoshka using Hi-C data from human IMR90 fibroblasts and mouse embryonic stem cells provided by Dixon et al. (1). The resulting interactions matrices were created with a bin size of 40kb and normalized for biases using an integrated probabilistic model (23). We applied our algorithm considering the *γ* values {0,0.05,0.1,0.15,…, 1}. For the IMR90 data, we used CTCF sites from (24) and for mESC from (25). Datasets for histone modification sites were obtained from various studies. For IMR90 we present results for H3K4me3 and H3K9me3 for which data is publicly available (26). Similarly, for mESC we present H3K4me3, H3K27ac (25) and H3K36me3 (27). Where needed, the relevant data was shifted to assemblies hg18 and mm9 for human and mouse data, respectively, using the UCSC liftover tool (28).

Relevant results from the mouse embryonic stem cell data are compared against the hierarchy generated by TADtree (19). We choose to use their result where, on average, 1.6 domains are allowed per megabase on each chromosome, since this returns the closest number of domains to our results.

Similarly, we also compare conservation and enrichment results against randomized hierarchies which are generated such that the following features of the hierarchy are preserved while shuffling the order of domains and non-domains:

1. The number of domains at each level of the hierarchy.
2. Sizes of the domains, as well as the regions between the domains.
3. The structure of the hierarchy, such that the nesting of the domains is preserved and sub-domains shuffle within the shuffled super-domain.

## RESULTS

### CTCF enrichment at domain boundaries

It has been shown that the protein CTCF binds many known insulator or barrier elements in the genome which tend to lie at the borders of chromatin domains (29). Hence, enrichment of CTCF binding has been used as a measure of quality of domains predicted by previous works (1, 15). We show that our hierarchical domains are highly enriched for CTCF at their boundaries for both the human and mouse datasets (see Figure 2a, b). We also compare against the mouse domains given by TADtree (19) and show that our domains appear more enriched with a sharper peak around the domain boundaries (see Figure 2c). This suggests that our hierarchical domains are more closely linked with biologically functional sites in the genome. This is also shown by the depletion of CTCF towards the center of the domains.

**Figure 2.**
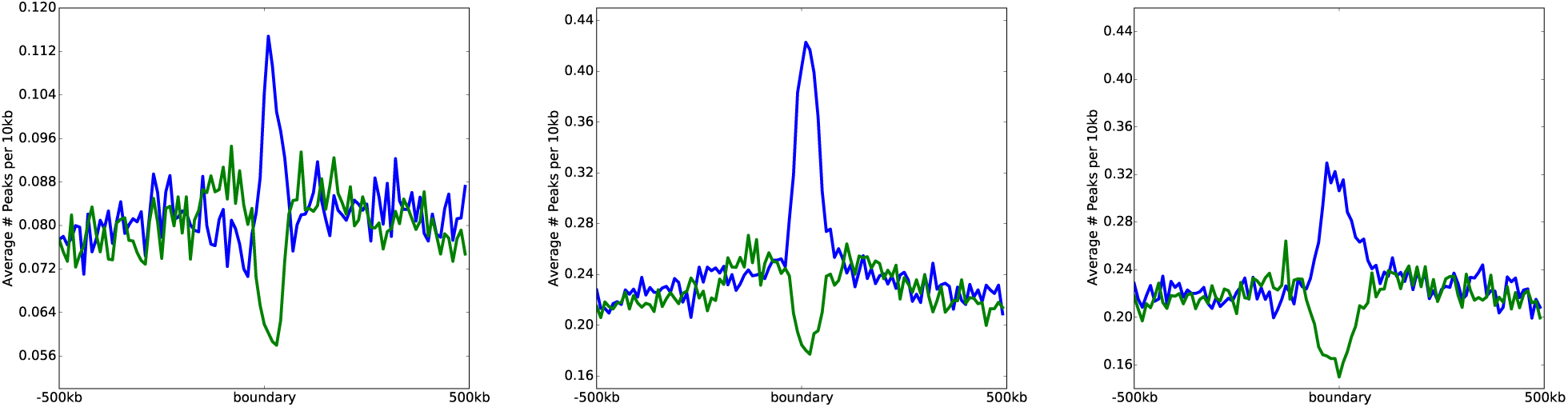
Enrichment of CTCF in domains predicted in both human and mouse data using Matryoshka. For mESC data, enrichment results are also provided for TADtree. The blue line shows the average number of CTCF peaks (averaged over 10kb intervals) centered at the boundaries we predict and 500kb on each side of these boundaries. Similarly the green line shows the average number of peaks 500kb on each side of the center of the predicted topological domains.

Apart from these, we also compared the ratio of CTCF sites overlapping with our domain boundaries in the IMR90 dataset against 1000 randomized hierarchies, generated as explained above. For the whole genome, the 1000 randomized hierarchies had a much lower ratio (p-value 0.001). We repeated the same analysis for each level of the hierarchy. For the first 4 levels, all the randomized domains have a lower ratio (p-value 0.001); for level 5, however, the result we obtain is not statistically significant (p-value ≤ 0.24). The higher values at this level may be due to the much smaller number of domains, combined with the decreasing sizes along the hierarchy. These tests act as a control to show that the enrichment results we observe at multiple levels of our hierarchy are unlikely to occur by chance.

### Histone modification analysis at domain boundaries

Histone modification marks are also used for analysis of chromatin domain boundaries since many of these are known to coincide with regions of enhancer-promoter interactions, resulting in active transcription (5, 30). For the hierarchical domains predicted in the human IMR90 dataset, we show they are enriched for H3K4me3 (see Figure 3a). These factors are associated with promoters in the mammalian cells (31). In contrast, there is a depletion of H3K9me3 marks at the boundaries, which are not associated with promoters as predicted by Dixon et al. (1) (see Figure 3b). In a similar way, we analyzed the hierarchical domains predicted by our algorithm and compared them against those predicted by TADtree. We show enrichment for several histone modifications and a higher average number of peaks within close proximity of our domain boundaries, as compared against those from TADtree. We analyzed H3K4me3 marks, which are indicative of active promoters in mice (25); H3K27ac marks, known to be associated with active enhancers (and therefore abundant around boundaries and within domains) (32) and H3K36me3 marks that are linked with actively transcribed genes (31) as well as promoter clusters (30). These have been shown to be enriched around domain boundaries predicted by earlier tools (1, 15) and we therefore use these to analyze quality of our hierarchical domain boundaries.

**Figure 3.**
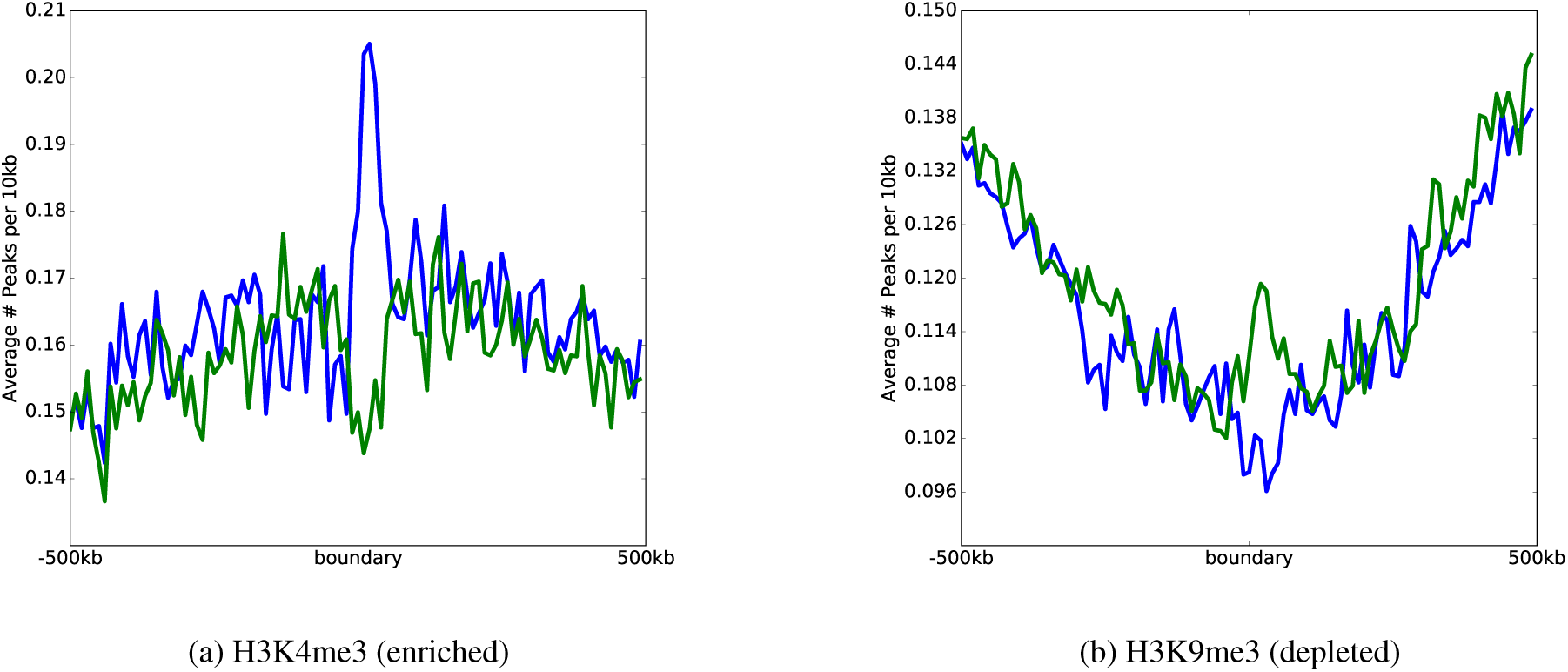
Histone modification analysis in Matryoshka domains from IMR90 data. The blue lines are for boundaries and green for midpoints of topological domains, as explained previously.

**Figure 4.**
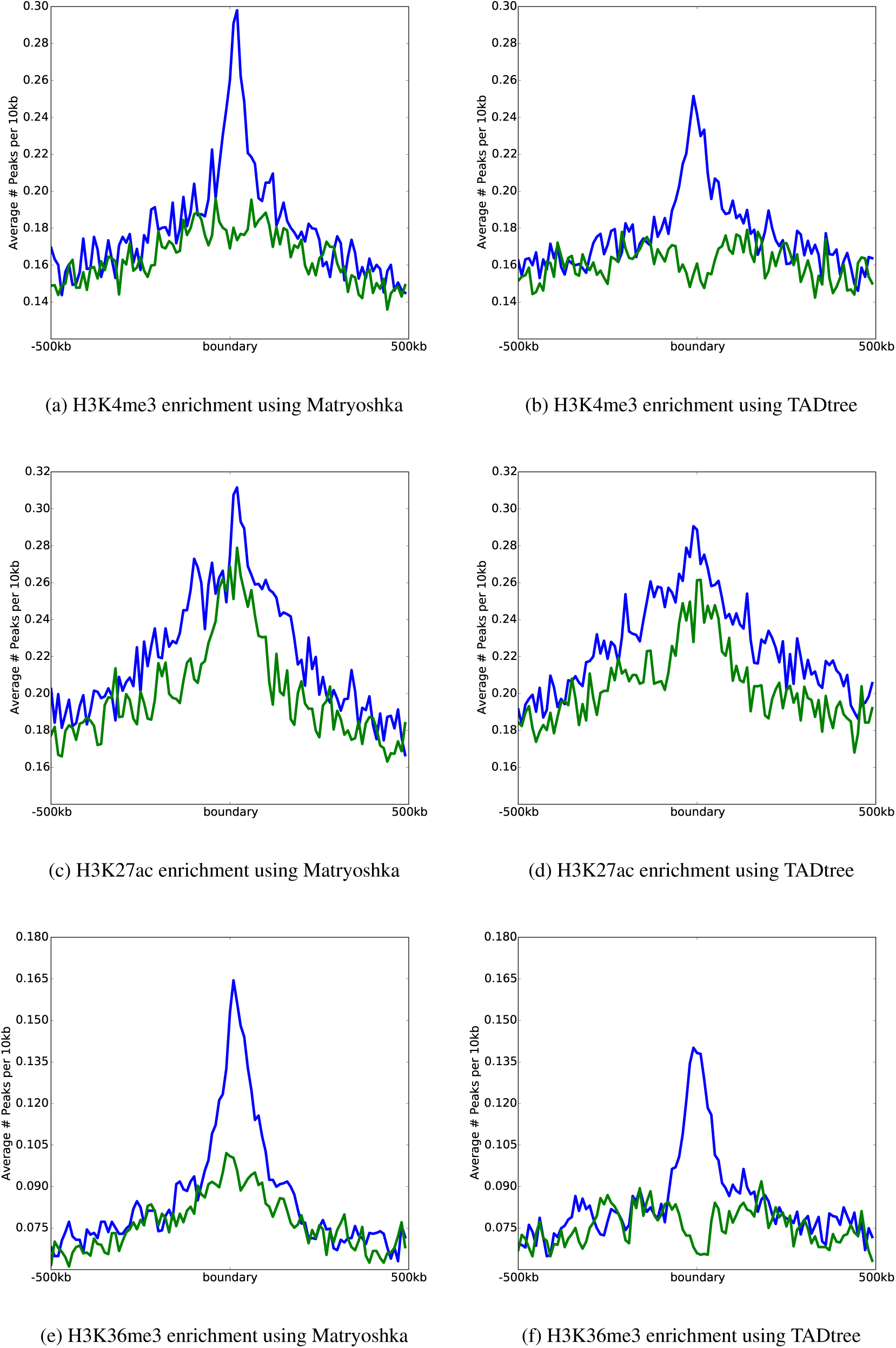
Histone modification analysis in domains from mESC data compared against domains from TADtree showing higher enrichment around domain boundaries predicted by Matryoshka. The blue lines are for boundaries and green for midpoints of topological domains, as explained previously.

### Conservation of hierarchy across species and cell types

Previous studies have shown that chromatin domains are conserved across species and cell types (1). We show that not just the domains we predict, but the hierarchical structure is conserved at each level as well. We do so by comparing domains in the whole genome, as well as those at individual levels of the hierarchy. Since we are comparing human fibroblast data with mouse embryonic stem cell, results would reflect conservation across both species and cell type. In order to compare the datasets, we use the UCSC liftover tool to convert domains from one set to the other. We calculate the ratio of overlap between two sets of domains, *D_i_* and *D_j_*, as follows:

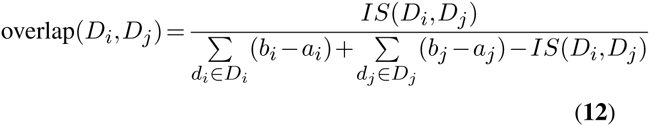
where *d_i_ =* [*a_i_,b_i_*] and IS is simply the sum of the lengths of intersecting regions from the two sets, 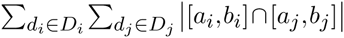.

We converted domains from the IMR90 data to the mESC data and calculated the overlap between this new domain set and the domains from mESC. Similarly, we converted the mouse domains and compared against human domains. As control, we randomized the hierarchical domains predicted by our program and repeated the procedure on these randomized sets. The randomized results presented are an average over a 1000 randomized domain trials. These results are presented in Table 1. Overall, we see a greater overlap between predicted domains in the whole genome as compared to randomized domains, showing that they are conserved across the datasets. This is also true for the higher (i.e. coarser) levels of the hierarchy. The discrepancy at lower levels could reflect that superdomains are conserved, whereas the subdomains allow for the variation across cell types (33). It has been predicted that larger domains are stable across cells and changes at a smaller level correspond to differentiation and variation in gene expression (16). These results reflect the biological significance of chromatin structure and domains that have been conserved across evolution and are an important property of the genomic architecture. It is also possible, of course, that lack of statistically-significant conservation at lower (i.e. finer) levels of the hierarchy results from the inevitable loss of data when lifting-over between species and cell types, and of the resolution limits of the original data.

**Table 1.**
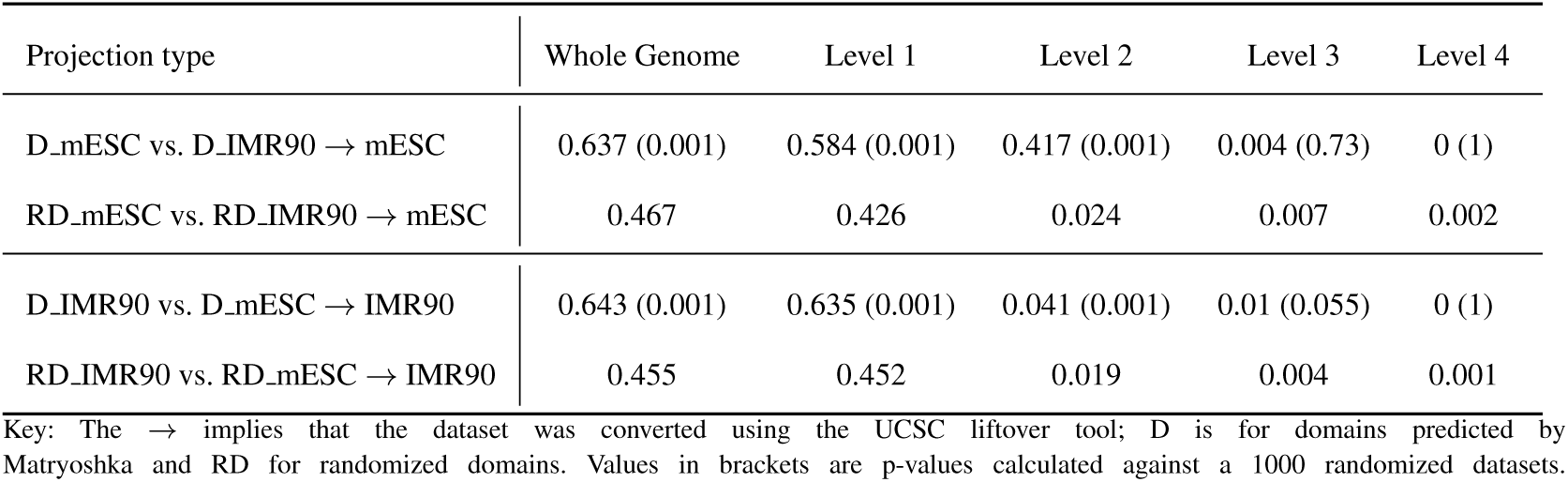
Conservation of domain structure across species and cell types, as measured by overlap as specified in equation **12**.

### Overlap with interacting regulatory elements

Enhancers and promoters regulate gene expression but can frequently be at long distances from the genes that they control. It is predicted that associated enhancers and promoters are more likely to interact within the same topological domain as compared to across domains (25). To test this for our domains, we used enhancer-promoter units (EPUs) in mESC defined by (25) and enhancer-promoter pairs in IMR90 predicted by (34), lifted over to hg18. We compared the ratio of these clusters or pairs that are completely nested within domains predicted by our algorithm against 1000 randomized hierarchies. For the mouse dataset, we find that 56% of the EPUs are completely nested within the domains predicted, compared to an average of 26% in the randomized domains (p-value 0.001). Similarly, for the domains in IMR90, 43% of the enhancer-promoter pairs were nested, compared to an average of 27% in the random selection (p-value ≤ 0.06). These show a strong correlation between topological domains and regulatory elements in the genome, with more interaction within a domain and relatively greater insulation across domains. These also reflect the functional role that topological domains play in gene expression and regulation. Further analysis is required to study the functional and biological relationship between genes from the same domain and effects on their expression with changes in chromatin architecture and domain nesting.

### Runtime analysis

Matryoshka, using *γ* values from 0 to 1 (inclusive) with a step size of 0.05, takes only 1-2 minutes to run on 40kb resolution Hi-C data from human chromosome 10. For human fibroblast, processing data for 22 chromosomes in total, took 39 minutes and 42 seconds, on a personal computer with 1.8GHz Intel Core i7 and 8Gb of RAM. On the same computer, the mouse embryonic stem cell data took 31 minutes and 11 seconds for the entire dataset, containing 19 chromosomes. In comparison with this, TADtree can take several hours to run on data from a single chromosome depending on the choice of parameters by the user. Hence, our method provides an efficient way for predicting hierarchical chromatin structure using Hi-C data.

## DISCUSSION

Analysis of chromatin conformation data has revealed the hierarchical nature of chromatin folding but there are no efficient tools that allow the extraction of this hierarchy from raw chromatin conformation capture data. In this paper, we presented a tool, Matryoshka, that predicts the nested structure of chromatin domains from raw Hi-C interaction matrices. Domains are extracted independently across a wide range of different scales using a variant of the method of Filippova et al. (15). Subsequently, our method effectively predicts the number of levels for the hierarchy based on the variation among domains at multiple resolutions. The distance metric used for clustering reflects the variation in domains at different resolutions and therefore a greater variation implies a larger number of possible nested domains. The algorithm is completely data-driven, and the only input required from the user is the maximum *γ* value, for which an appropriate value can be set based on properties of the input data. We show that the domain boundaries predicted by Matryoshka are highly enriched for insulator and barrier-like elements. The role of these elements in gene regulation and their relationship with chromatin domains has been previously validated.

Further, we show the relationship between hierarchical domains in mouse and human data and demonstrate that superdomains (the coarse-grained levels of our hierarchy) are conserved. A more extensive study of the complete structure across various species, and not just the set of linear domains, could contribute to our understanding of the evolution of DNA structure. It would help to analyze how gene regulatory mechanisms vary at the super and subdomain levels in different organisms. Previous studies have predicted that stable larger domains may have a role to play in cell-cycle regulation and timing, whereas changes within these domains could control gene expression and differentiation (35, 36). Our method provides an efficient way to classify the structures of domains at different scales, enabling us to compare them across cell types.

Similarly, the role of hierarchical domains in diseased cells could be analyzed. It is known, for example, that chromatin structure is correlated with the activity of cancerous cells (3, 37, 38). A comparison of the nested structure in diseased and normal cells could give insights into the regulatory methods employed by healthy cells and how these are perturbed in the disease state. Further analysis would be required to determine if differences are in superdomains or subdomains, and the functions to which the genes in these domains correspond. Our algorithm allows for these studies to be carried out efficiently on a large number of datasets.

Apart from these investigations, future work on chromatin structure using higher resolution data would give more insights into how domain hierarchies vary at a finer level and the significance of nested domains may become be more evident. Combining chromatin structure data with other sequencing assays is an interesting direction to explore and will enable us to relate variation in expression levels with topological domains (39, 40). The relationship between nesting of domains and differential expression can be be studied in a similar way. The importance of the 3-dimensional structure of chromatin may only become fully apparent when analyzed in conjunction with other assays, so that we can explore how changes in chromatin architecture correlate with other functional changes in the cell.

